# β-barrel proteins dictate the effect of core oligosaccharide composition on outer membrane mechanics

**DOI:** 10.1101/2024.09.02.610904

**Authors:** Dylan Fitzmaurice, Anthony Amador, Tahj Starr, Glen M. Hocky, Enrique R. Rojas

## Abstract

The outer membrane is the defining structure of Gram-negative bacteria. We previously demonstrated that it is critical for the mechanical integrity of the cell envelope and therefore to the robustness of the bacterial cell as a whole. Here, to determine the key molecules and moieties within the outer membrane that underlie its contribution to cell envelope mechanics, we measured cell-envelope stiffness across several sets of mutants with altered outer-membrane sugar content, protein content, and electric charge. To decouple outer membrane stiffness from total cell envelope stiffness, we developed a novel microfluidics-based “osmotic force extension” assay. In tandem, we developed a simple method to increase throughput of microfluidics experiments by performing them on color-coded pools of mutants. Using *Escherichia coli* as a model Gram-negative bacterium, we found that truncating the core oligosaccharide, deleting the β-barrel protein OmpA, or deleting lipoprotein outer membrane-cell wall linkers all had the same modest, convergent effect on total cell-envelope stiffness but had large, varying effects on the ability of the cell wall to transfer tension to the outer membrane during large hyperosmotic shocks. Surprisingly, altering lipid A charge had little effect on the mechanical properties of the envelope. Importantly, the presence or absence of OmpA determined whether truncating the core oligosaccharide decreased or increased envelope stiffness (respectively), revealing sign epistasis between these components. Based on these data we propose a specific structural model in which the chemical interactions between lipopolysaccharides, β-barrel proteins, and phospholipids coordinately determine cell envelope stiffness, and the ability of the outer membrane to functionally share mechanical loads with the cell wall.

**Statement of Significance:** The outer membrane is the defining cellular structure of Gram-negative bacteria, a group that contains many important pathogens like *Escherichia coli*, *Vibrio cholerae*, and *Pseudomonas aeruginosa*. One role of the outer membrane is to block the uptake of small molecules like antibiotics. However, it is becoming increasingly clear that it also functions as a structural exoskeleton that is critical for the cell’s ability to cope with internal and external mechanical forces. Here, we carefully dissect the molecular basis for the load-bearing capacity of the outer membrane by screening a set of mutants with a new cell biophysics assay.

## Introduction

The cell envelope of Gram-negative bacteria (Fig. 1A) is a permeability barrier and exoskeleton that mediates all interactions between the bacterial cell and its environment, defines cell shape, and confers robust mechanical properties to the cell. This latter function is vital to bacteria during osmotic fluctuations^1^, growth in confined spaces^2^, and antibiotic exposure^3^. The envelope is comprised of three essential layers: the plasma membrane, the peptidoglycan cell wall, and the outer membrane - an atypical bilayer with a phospholipid inner leaflet and an outer leaflet composed of complex macromolecules called lipopolysaccharides (Fig 1B). Until recently, the robust mechanical properties of the cell envelope were exclusively attributed to the covalently cross-linked cell wall^4, 5^. However, we demonstrated that the outer membrane of *Escherichia coli* is actually stiffer than its cell wall with respect to tension in the cell envelope^1^. Furthermore, several major genetic and chemical perturbations to the outer membrane dramatically reduced its ability to bear mechanical forces, leading to fragile cells. There are three immediate questions motivated by this discovery: first, with respect to osmotic variation (during which the outer membrane is mechanically engaged), what are the constitutive mechanical properties of the outer membrane (*e.g.*, linear versus non-linear)? Second, are there key molecules or moieties that determine these properties, or do they emerge from the outer membrane complex as a whole? Third, is there a specific architecture underlying how outer membrane components are connected that allows them to bear mechanical loads (e.g. “in series” or “in parallel”)?

**Figure 1.**
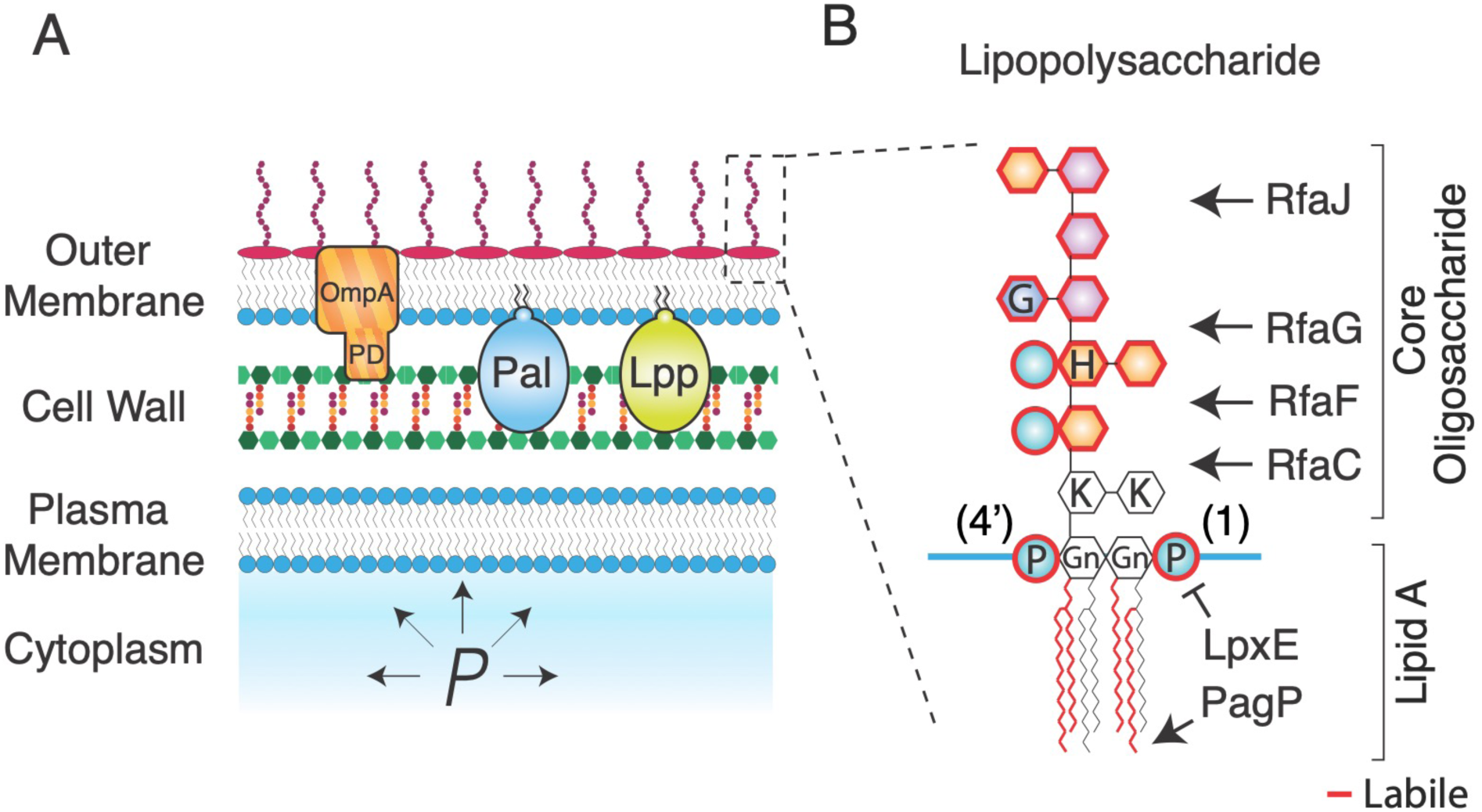
The Gram-negative cell envelope is complex. A) Schematic of the Gram-negative cell envelope. PD: periplasmic domain of OmpA. *P*: turgor pressure. B) Chemical structure of lipopolysaccharide with structure-modifying enzymes. Moieties outlined in red are enzymatically labile. Gn: glucosamine, P: phosphate, H: heptose, G: glucose, K: keto-deoxyoctulosonate.

The intermolecular ionic bonds between the lipid A domains of lipopolysaccharides (Fig. 1B) are, collectively, a leading candidate for a mechanical chassis within the outer membrane. Lipid A consists of a glucosamine disaccharide head group linked to six acyl chains that interface with the inner leaflet of the outer membrane. The head group is phosphorylated at the 1 and 4’ carbons, which allows lipopolysaccharides to bind to one another in the presence of divalent magnesium ions, which mediate intermolecular ionic “salt bridges” between phosphate groups (Fig. 1B). Given that the outer membrane is not fluid (proteins do not diffuse within it^6–8^), it is likely that these ionic bonds create a solid lipopolysaccharide-magnesium gel. Chelation of magnesium away from the outer membrane by ethylenediaminetetraacetic acid (EDTA) results in a porous, weak outer membrane as would be expected if the salt bridges were key load-bearing bonds^1^. However, it is likely that magnesium chelation completely destabilizes the outer membrane, making it difficult to decouple the mechanical contributions of the salt bridges from those of other interactions that are also eliminated upon EDTA treatment. For example, forces could also be borne by hydrophobic interactions between the acyl moieties of lipid A, which are indirectly disrupted by EDTA.

Bacteria can enzymatically modify lipid A, including its electrical charge, in response to environmental stimuli. For example, in response to weak acids *E. coli* adds phosphoethanolamine and 4-aminoarabinose to the lipid A phosphates, which makes it less anionic and results in increased outer membrane permeability^9^. In principle, such modifications could also provide a way for bacteria to adaptively modulate their mechanical properties if these properties were dependent, for example, on electric charge density. Importantly, by ectopically expressing the specific enzymes that modify lipid A and alter its charge it is possible to controllably study the effect of these modifications^10^.

Besides lipid A, lipopolysaccharide features two polysaccharide moieties, which could also provide mechanical contributions to the outer membrane. The core oligosaccharide is a 10-residue heteropolymer (Fig. 1B) that is usually conserved within genera or families of Gram-negative bacteria^11^. Core oligosaccharide synthesis occurs sequentially by the Rfa monosaccharide transferases, such that the deletion of one of these enzymes results in an oligosaccharide that is truncated at the residue attached by that enzyme (Fig. 1B). Undomesticated wild-type Gram-negative bacteria ligate an additional polysaccharide called the O-antigen to the terminal core-oligosaccharide residue; the composition of the O-antigen is highly variable across bacterial species and strains. Interestingly, the O-antigen can increase mechanical integrity to the cell even if it is electrically neutral^1^. Whether the specific length or composition of the core oligosaccharide also affects outer membrane stiffness is unknown, however certain truncation mutants (△*rfaC* and △*rfaG*; Fig. 1B) result in increased outer membrane vesiculation^12^, which may point to a weakened outer membrane.

In addition to lipopolysaccharides, the outer membrane is densely loaded with proteins^13^. Most of these are β-barrel proteins: β-sheets that are folded into transmembrane cylinders by the Bam complex^14, 15^. Certain β-barrel proteins are selective molecular pores (“porins”). For example, OmpF and OmpC are highly abundant β-barrel porins in *E. coli* whose expression is coordinately regulated in response to extracellular osmolarity, while LamB is specifically expressed to mediate uptake of maltose^16, 17^. Whether these porins are also important for the mechanical integrity of the cell envelope is unknown. However, the structure and folding pattern of the β-barrel protein EspP is sensitive to tension in the outer membrane, providing insight into how these proteins could bear mechanical forces^18^.

One protein known to be critical for the mechanical integrity of the cell envelope is the highly abundant β-barrel protein OmpA: the deletion of this protein causes drastic weakening of the outer membrane^1^. However, as for the case of EDTA it is unknown if this means that OmpA is a specific mechanical element or if its elimination causes global destabilization of the cell envelope. This question in particularly relevant to the case of OmpA since, unlike other β-barrel proteins, it possesses a periplasmic domain that specifically binds to the peptidoglycan cell wall, making it likely that its deletion has pleiotropic effects on the global stiffness of the cell envelope.

Indeed, OmpA is one of three proteins that connect the outer membrane to the cell wall (Fig. 1A). The other two key linkers, Pal and Lpp, are lipoproteins. Like OmpA, Pal non-covalently binds the cell wall and is critical for mediating constriction of the outer membrane during cell division as part of the Tol-Pal complex^19, 20^. In contrast, the lipoprotein Lpp is covalently ligated to the cell wall and acts as a molecular pillar that determines the width of the periplasm^21, 22^. Collectively, OmpA, Pal, and Lpp prevent loss of outer membrane material via vesiculation^23^. We previously found that bacterial mutants that lack any of these proteins have a weaker cell envelope and are highly susceptible to lysis upon repeated osmotic shocks^1^. It is unknown, however, if these phenotypes are due intrinsic load-bearing capacity of the proteins themselves or due to the de-coupling of the outer membrane and cell wall that results from their deletion. Furthermore, it is unknown whether during modest osmotic shocks these molecular staples are important for transferring mechanical forces between the cell wall and outer membrane.

Our previous assays for interrogating cell envelope mechanics were useful for highlighting the critical contribution of the outer membrane to total cell-envelope stiffness, but were limited due to issues of specificity, precision, and throughput. One key assay we developed was a microfluidics “plasmolysis-lysis” experiment that measured the ratio between the stiffnesses of the outer membrane, *k*_om_, and that of the cell wall, *k*_cw_^1^ (Fig. S1). In this assay, cells were subjected to a large (3 M) hyperosmotic shock and subsequently perfused with detergent, which caused cell lysis and dissolved the outer membrane. Although turgor pressure was completely depleted, we found that after the hyperosmotic shock (but before detergent perfusion) the cell wall was still stretched because of its association with the outer membrane, which prevented the wall from relaxing to its rest state by bearing in-plane compressive stress (Fig. S1). By quantifying the contractions of the cell wall upon hyperosmotic shock and lysis and treating the outer membrane and cell wall as parallel linear springs, we estimated the ratio *k_om_/k_cw_* (Eq. 1, *Methods*). For *E. coli*, we found that the outer membrane stiffness was about 1.5 times that of the cell wall, which pointed to the potential importance of the outer membrane as a mechanical element. Furthermore, mutations that reduced this ratio sensitized bacteria to osmotic fluctuations. Collectively, this pipeline provided a useful empirical quantification of cell-envelope mechanical properties. However, it did not alone decouple the stiffness of the outer membrane from that of the cell wall, which is particularly important for assessing pleiotropy.

An important aspect of the plasmolysis-lysis assay is that for mutants with impaired connections between the outer membrane and cell wall, the large hyperosmotic shock is likely to cause partial detachment of the outer membrane from the cell wall, and therefore during this treatment the outer membrane cannot share envelope tension with the cell wall, regardless of its intrinsic stiffness. Therefore, the quantity *k_om_/k_cw_* that this assay reports is more precisely the ratio between the “effective outer membrane stiffness” and the stiffness of the cell wall.

In another assay, we probed cell-envelope mechanics by measuring cell-envelope deformation in response to a small hyperosmotic shock of a single magnitude (*ΔC*=200 mM). This caused a defined reduction in turgor pressure (*ΔP = -RTΔC*), where *R* is the ideal gas constant and *T* is the temperature) that partially deflated the cell. We demonstrated that the degree of this deformation was related to the mechanical properties of the cell envelope. However, using a single shock magnitude did not provide the specific scaling relationship between envelope deformation and pressure changes (*e.g.* linear vs. non-linear). This information is important since the cell wall exhibits non-linear strain-stiffening as measured via atomic force spectroscopy^24^; if this behavior was also occurring during osmotic shocks it would obscure the meaning of deformation at a single shock magnitude.

Finally, all existing methodologies to measure cell-envelope mechanical properties at the single-cell level - including atomic force microscopy^24^ and cell bending assays^25^ - are inherently low throughput, typically requiring several replicate experiments for each bacterial strain or mutant. This limits our ability to efficiently screen enough mutants to obtain a comprehensive understanding of the relationship between cell envelope composition and cell-envelope mechanical properties.

In sum, due to technical limitations we lack a precise understanding of the constitutive mechanical properties of the cell envelope, and the molecular components that confer its mechanical integrity. To address this, we developed a new “osmotic force-extension assay” to quantitatively measure cell-envelope stiffness with more precision than previous assays were capable of (Fig. 2A-D). Using this assay, we found that the cell envelope is a linear elastic material with respect to in-plane compression. In combination with the plasmolysis-lysis assay (Fig. S1), the osmotic-force-extension assay also allowed us to de-couple effective outer membrane stiffness from cell-wall stiffness. To accelerate throughput, we developed a method to color-code bacterial strains using combinations of non-toxic fluorophores, which allowed us to perform our microfluidics assays on pools of mutant bacteria (Fig. 2E). Using these assays, we systematically measured how genetic alterations of three families of molecules and moieties within the outer membrane – core oligosaccharides, β-barrel proteins, and lipid A – affect cell-envelope and outer membrane stiffness. A simple but important result of this analysis was that major perturbations to any of these components had the same quantitative effect on cell envelope stiffness, indicating that this property arises from the collective assembly of envelope components. We also found that while systematic truncation of the core oligosaccharide of *E. coli* monotonically decreased cell-envelope and outer-membrane stiffness, the same mutations monotonically increased these properties in the absence of OmpA. Based on these results, we propose a model for how the interactions between core oligosaccharides, β-barrel proteins, and phospholipids coordinately determine the mechanical integrity of the outer membrane. Collectively, our analysis provides a more highly-resolved picture of the mechanical infrastructure of the cell envelope than was possible with previous methods, and provides new broadly useful assays for interrogating this infrastructure.

**Figure 2.**
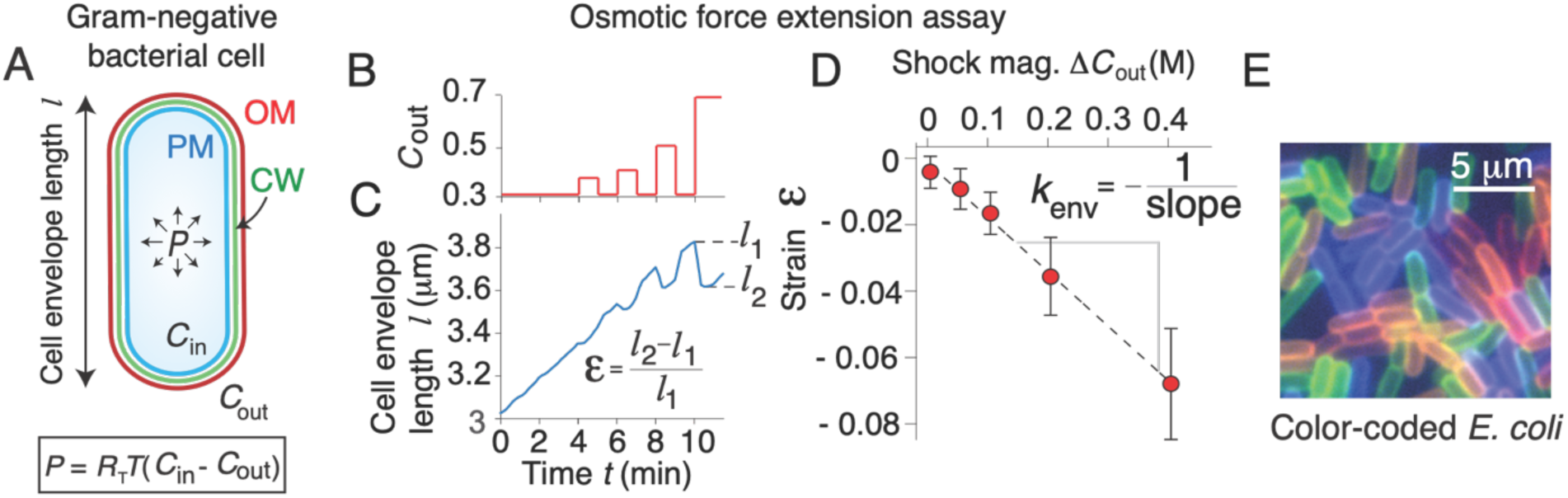
An osmotic force extension assay measures total cell-envelope stiffness. A) (top) Diagram of a Gram-negative bacterial cell inflated with turgor pressure, *P*. OM: outer membrane, CW: cell wall, PM: plasma membrane, *C*_in_: cytosolic osmolarity, *C*_out_: osmolarity of the growth medium. (bottom) Turgor pressure is proportional to the difference between the cytosolic and growth medium osmolarities, where *R*_T_ is the gas constant and *T* is the temperature. B) Osmolarity of growth medium versus time during an osmotic-force extension experiment. C) Cell-envelope length during an osmotic-force extension experiment. D) Mechanical strain in cell length versus shock magnitude. The dotted line is the best fit using linear regression. Cell envelope stiffness, *k*_env_, is calculated as the inverse of the slope of the regression. E) A pool of color-coded *E. coli* cells.

## Results

### An osmotic force extension assay precisely measures cell envelope stiffness

Our goals were to develop a precise and efficient method for measuring the constitutive mechanical properties of the cell envelope, to decouple the stiffness of the outer membrane from the cell wall, and to apply these methods to a wide range of genetic mutations (and combinations thereof) to dissect the mechanical structure of the cell envelope. To begin, we developed a new “osmotic force extension” assay (Fig. 2A-D) in which we subjected cells to a series of hyperosmotic shocks of increasing magnitude, and measured the resulting contractions of the cell envelope (strain in cell length) caused by each shock (Fig 2B,C). We discovered that the dependence of strain on shock magnitude was precisely linear for shocks up to 400 mM, which allowed us to empirically define cell envelope stiffness, *k*_env_, as the inverse of the slope of this dependence (Fig. 2D). This result validated the treatment of the cell wall and outer membrane as linear springs in the plasmolysis-lysis assay. Therefore, by combining the two assays we could empirically solve for the stiffnesses of the cell wall and outer membrane in terms of experimentally measurable quantities (*Methods*).

In concert with this analysis, to accelerate the throughput of our experimental pipeline we invented a method to color-code bacterial strains with combinations of non-toxic fluorophores (Fig. 2E). This allowed us to pool up to 9 color-coded mutants and perform our microscopy/microfluidics assays on the entire pool at once, greatly increasing the throughput of our mechanical characterization. An additional benefit of this method is that we could include the isogenic wild-type background in each pool of mutants, thereby providing an internal control in all experiments.

### Cell envelope stiffness is correlated with core oligosaccharide length

We first measured the effect of truncations of the core oligosaccharide on the mechanical properties of the *E. coli* outer membrane. Because of the large contribution of the outer membrane to envelope stiffness, we hypothesized that even minor alterations to the core oligosaccharide would meaningfully affect total envelope stiffness. When we applied the methods described above to a set of mutants with deletions of the *rfa* genes, we found that total envelope stiffness was strongly correlated with core oligosaccharide length across two wild-type backgrounds of *E. coli* (MG1655 and BW25113; Fig. 3A). Furthermore, this dependence arose directly from weakening of the outer membrane (Fig. 3B), whereas the stiffness of the cell wall did not systematically depend on core oligosaccharide length (Fig. 3C). Complete removal of the “outer core” (by deletion of *rfaC*) leaving only the essential “inner core,” resulted in a 72% reduction in outer membrane stiffness (Fig. 3B).

**Figure 3.**
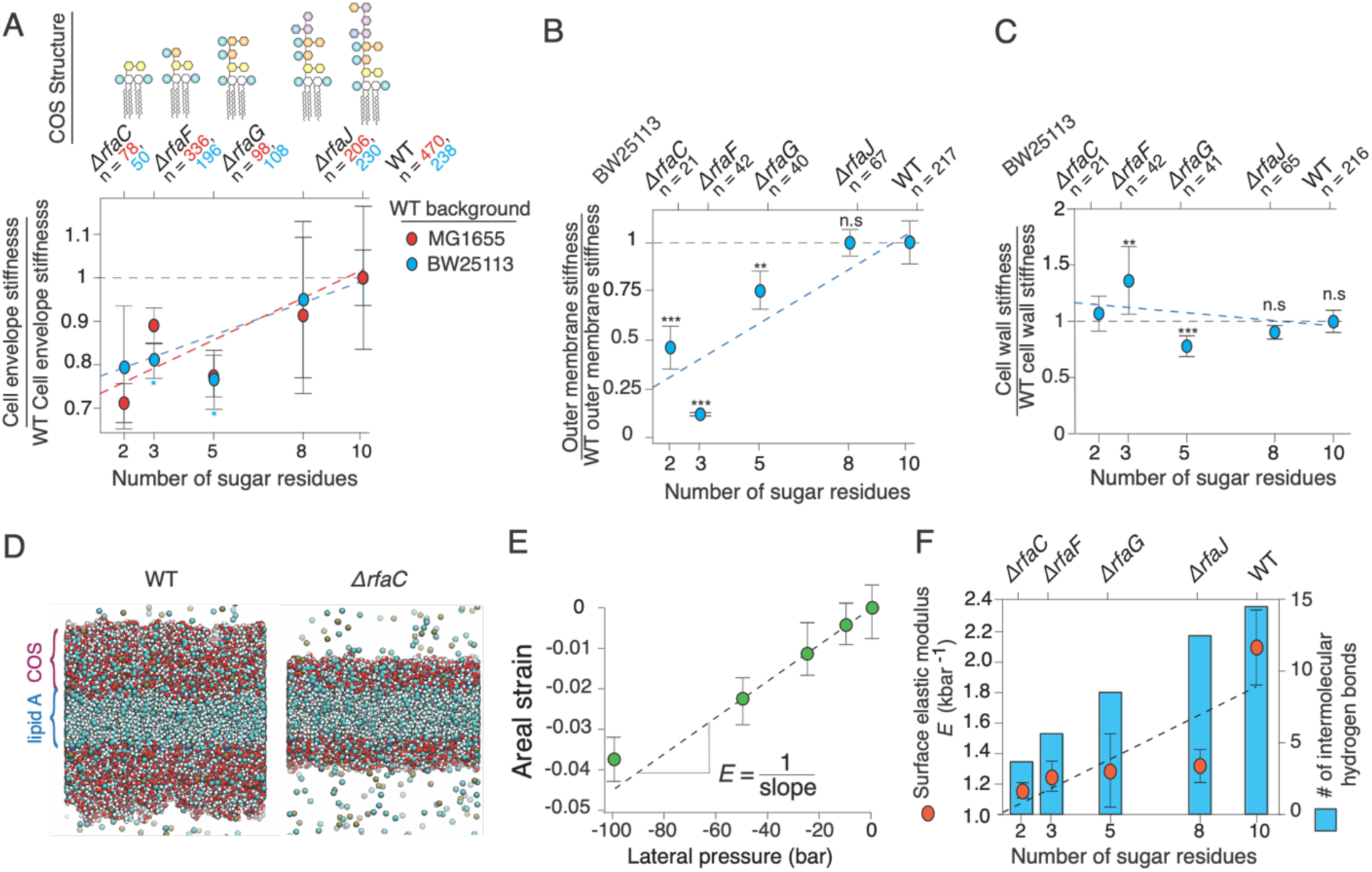
Cell envelope and outer membrane stiffness are proportional to lipopolysaccharide length. A) Cell envelope stiffness versus core oligosaccharide length, normalized by wild-type cell-envelope stiffness; *n* = 50, 196, 108, 230, 238 for Δ*rfaC*, Δ*rfaF*, Δ*rfaG,* Δ*rfaJ* and BW25113 wild-type cells, respectively. Error bars indicate +/- 1 s.d. across 2-3 experiments per mutant. C) Outer membrane stiffness versus core oligosaccharide length, normalized to wild-type outer membrane stiffness; *n* = 21, 43, 41, 78, 220 for Δ*rfaC*, Δ*rfaF*, Δ*rfaG,* Δ*rfaJ* and BW25113 wild-type cells, respectively. Most of the core *rfa* mutants in the MG1655 background lysed when we performed the plasmolysis-lysis assay on them and therefore we could not decouple outer membrane and cell wall stiffness. (D-F) are results from MD simulations. D) Illustration of simulated wild-type lipopolysaccharide bilayer (left) and Δ*rfac* lipopolysaccharide bilayer (right). E) Areal strain versus lateral pressure for the wild-type simulated lipopolysaccharide bilayer. F) (orange circles, left axis) Surface (2-D) elastic modulus versus core oligosaccharide length for simulated lipopolysaccharide bilayers. Error bars indicate +/- 1 s.d. across three 20 ns time-windows after the system had reached its maximum contraction. (blue bars, right axis) Mean number of intermolecular hydrogen bonds at a given time versus core oligosaccharide length.

To explore the origin of the forces borne by the core oligosaccharides, we used all-atom molecular dynamics simulations to test the effect of outer core oligosaccharide truncations on the stiffness of the outer membrane *in silico*. In molecular dynamics simulations, we subjected model membranes to lateral (in-plane) compressive tension characteristic of the tension experienced by the outer membrane during our experiments (Fig. 3D). We found that the dependence of areal compression on lateral pressure was approximately linear for compressions up to 5% (Fig. 3E) and thus we identified the surface elastic modulus, *E*, as the inverse of the slope of this dependence. By considering our results from molecular dynamics simulations in concert with our experimental values of outer membrane stiffness and the structure of lipopolysaccharide (Fig. 1B), we inferred the types of intermolecular interactions between core oligosaccharides that bear mechanical forces during surface compression of the outer membrane. Specifically, we asked whether core oligosaccharides bear forces within ionic salt bridges between phosphate groups (mediated by divalent cations) or via hydrogen bonding between polysaccharides.

As in our experiments (Fig. 3A), the surface elastic modulus of the outer membranes *in silico* monotonically decreased as core oligosaccharide was truncated (Fig. 3F). The fully truncated core oligosaccharide resulted in a 43% reduction in simulated membrane stiffness compared to 72% measured experimentally.

In both experiment and simulation, deleting *rfaG*, which removes half of the sugar residues from the core oligosaccharide, resulted in a ≈25% decrease in outer membrane stiffness (Fig. 3A,F). Our molecular dynamics simulations demonstrated that this truncation removes approximately half of the intermolecular hydrogen bonds between core oligosaccharides (Fig. 3F). Since this mutation does not result in the removal of any phosphate groups, this analysis demonstrates that hydrogen bonds are important load-bearing bonds within the outer membrane.

Further truncations of the core oligosaccharide removed both sugar residues and phosphate groups. In our simulations, we explicitly confirmed the existence of salt bridges at phosphate groups by measuring the spatial distribution of divalent cation concentration across the direction normal to the bilayer – salt bridges appear as sharp peaks in this distribution (Fig. S2A). Experimentally, deleting *rfaF* and *rfaC*, which remove one phosphate group each (as well as 3 sugar residues total: 2 for *rfaF* and 1 for *rfaC*) resulted in large ≈25% decreases in relative outer membrane stiffness (Fig. 3A). This value was quantitatively similar to the effect of removing five sugar residues by deleting *rfaG*, suggesting that while the weaker hydrogen bonds collectively make a meaningful contribution to outer membrane stiffness, the stronger ionic bonds make a larger contribution on a bond-by-bond basis. However, contrary to our experimental data, deleting *rfaF* and *rfaC in silico* had a smaller effect on the stiffness of the outer leaflet bilayer.

### The mechanical contributions of the β-barrel and periplasmic domains of OmpA can be decoupled

There are several possible reasons for the partial discrepancy between our experimental and computational results, however a key difference between the actual outer membrane and our simulated outer membrane is that proteins are absent from the latter. In this light, we hypothesized that core oligosaccharide-protein interactions are important determinants of outer membrane stiffness. We tested this explicitly by first measuring the effect of deleting β-barrel proteins on outer membrane stiffness, and then testing the effect of these deletions in combination with truncations of the core oligosaccharide. We reasoned that non-additive effects of these combinations on stiffness would reveal genetic or structural interactions between core oligosaccharides and proteins.

We focused on OmpA because this also provided the opportunity of investigating the relative contributions of its β-barrel domain and the periplasmic domain (Fig. 1A) to the mechanical integrity of the envelope. When we subjected *τ1ompA* mutants from three wild-type backgrounds to the osmotic force extension assay, we found that this deletion resulted in a consistent ≈25% reduction in cell envelope stiffness (Fig. 4A). Interestingly, removing only the periplasmic domain had a quantitatively similar effect on envelope stiffness, suggesting that the periplasmic linker function rather than the β-barrel domain underlies OmpA’s mechanical contribution. However, consistent with our previous study^1^, we found that deletion of OmpA completely abolished the outer membrane’s contribution to envelope stiffness in the plasmolysis-lysis assay. As a result, the effective outer membrane stiffness we measured was close to zero (Fig. 4B). Interestingly, when we expressed only the linker-less β-barrel domain of OmpA, this partially restored outer membrane stiffness. While these data are consistent with the periplasmic linker being a key mechanical linchpin within the cell envelope, they also clearly demonstrate that the β-barrel itself plays a mechanical role in certain contexts.

**Figure 4.**
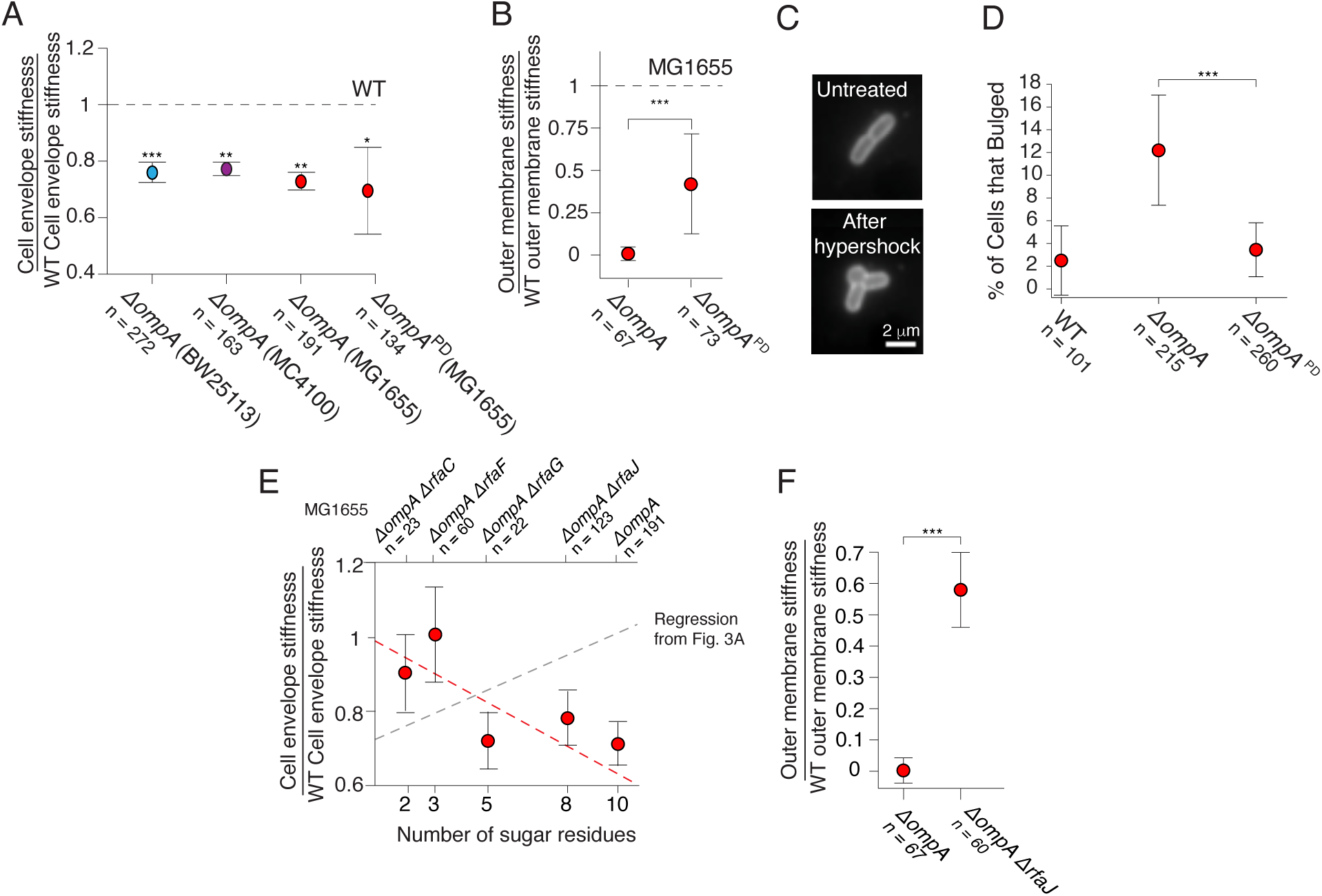
Mutations to *ompA* and *rfa* genes exhibit sign epistasis. A) Cell envelope stiffness for *ΔompA* mutants from three wild-type backgrounds, and for the deletion of the periplasmic domain (*ΔompA^PD^*) of OmpA in the MG1655 background. B) Outer membrane stiffness for the *ΔompA* and *ΔompA^PD^*mutants. C) A *ΔompA* cell before and after 400 mM hyperosmotic shock. D) Percentage of cells that developed outer membrane bulges after 400 mM hyperosmotic shocks. E) Cell envelope stiffness versus core oligosaccharide length for mutants in a *ΔompA* background, normalized by wild-type cell-envelope stiffness. F) Outer membrane stiffness for *ΔompA* and *ΔompA ΔrfaJ* mutants.

To further explore the contexts in which β-barrel or periplasmic linker were important, we labeled the outer membrane and explicitly measured its deformation during intermediate (400 mM) hyperosmotic shocks, which partially deplete pressure. We found that whereas the outer membrane of wild-type cells remained evenly attached to the cell wall, in a fraction of *ΔompA* mutant cells the hyperosmotic shock caused delamination of the outer membrane, leading to a large outer membrane bulge reminiscent of those observed after vancomycin treatment^26^ (Fig. 4C,D). Surprisingly, expressing the β-barrel domain of OmpA alone suppressed the bulging phenotype. That is, OmpA prevents delamination independent of its periplasmic linker.

Together, our data paint a complex picture of OmpA’s contribution to outer membrane mechanics. Deletion of the periplasmic linker is enough to modestly reduce envelope stiffness, to greatly reduce (effective) outer membrane stiffness, but presumably not enough to prevent Pal and Lpp from holding the outer membrane and cell wall together. However, additional deletion of the β-barrel domain is enough to loosen the attachment of the outer membrane to the cell wall and completely eliminate the effective contribution of the outer membrane to envelope mechanics during large hyperosmotic shocks (3 M) but not enough to weaken its contribution during modest ones (400 mM).

### Mutations to core oligosaccharides and OmpA exhibit sign epistasis

We next examined the genetic interactions between mutations to the core oligosaccharides and to OmpA. Surprisingly, we found that while truncating the core oligosaccharide predictably decreased cell-envelope and outer-membrane stiffness in the presence of OmpA (Fig. 3A), the same truncations increased cell envelope stiffness in the absence of OmpA (Fig. 4E).

Unfortunately, for all but one of the *ΔompA Δrfa* double mutants we were unable to perform our plasmolysis-lysis experiment due to cell lysis, and therefore we could not specifically decouple outer membrane stiffness for these strains. For the one double mutant that did survive (*ΔompA ΔrfaJ*, which possessed the smallest perturbation to the core oligosaccharide) we calculated that truncation of the core oligosaccharide greatly increased the contribution of the outer membrane to cell envelope stiffness compared to the single *ΔompA* deletion (Fig. 4F). In other words, mutations to *ompA* and the *rfa* genes result in what geneticists refer to as sign-epistasis, where the presence or absence of one gene determines the sign of the effect of a second gene on a given phenotype^27^. Further research is required to understand the molecular basis for this phenomenon, but we propose a simple putative model based on interactions between core oligosaccharides, β-barrel proteins, and phospholipids (*Discussion*).

### Mutants of outer membrane-cell wall linkers phenocopy *ΔompA* in envelope stiffness but not outer membrane stiffness

We next measured the effect of deletion of Lpp and Pal on cell envelope stiffness. Interestingly, we found that eliminating Lpp reduced total cell-envelope stiffness to precisely the same degree as eliminating OmpA or its periplasmic domain (Fig. 4A, 5A). Similarly, deletion of Lpp caused a reduction in outer membrane stiffness similar to that caused by deletion of OmpA’s periplasmic domain but less than the deletion of the entire OmpA protein (Fig. 4C, 5B). We propose that the precise quantitative correspondence between these mutations means that they are effectively leading to a convergent, modest structural collapse of the cell envelope that does not depend on the structure or copy number of the individual proteins. This likely means there is a threshold of outer membrane-cell wall connections that required to prevent this collapse. Deletion of Pal, however, had a stronger effect on cell envelope stiffness and a dramatic effect on effective outer membrane stiffness. In fact, when cells were treated with detergent after having been plasmolyzed (Fig. S1), the cell wall elongated instead of contracting, leading to negative values of outer membrane stiffness (Fig. 5B). The meaning of this is unclear, but one possibility is that in this mutant, the protoplast (plasma membrane and cytoplasm) can exert negative pressure on the cell envelope during plasmolysis, and because the outer membrane-cell wall links are severely undermined, the cell wall contracts below its rest length. In any case, it is clear from these measurements Pal is the most important of the three linkers mechanically.

**Figure 5.**
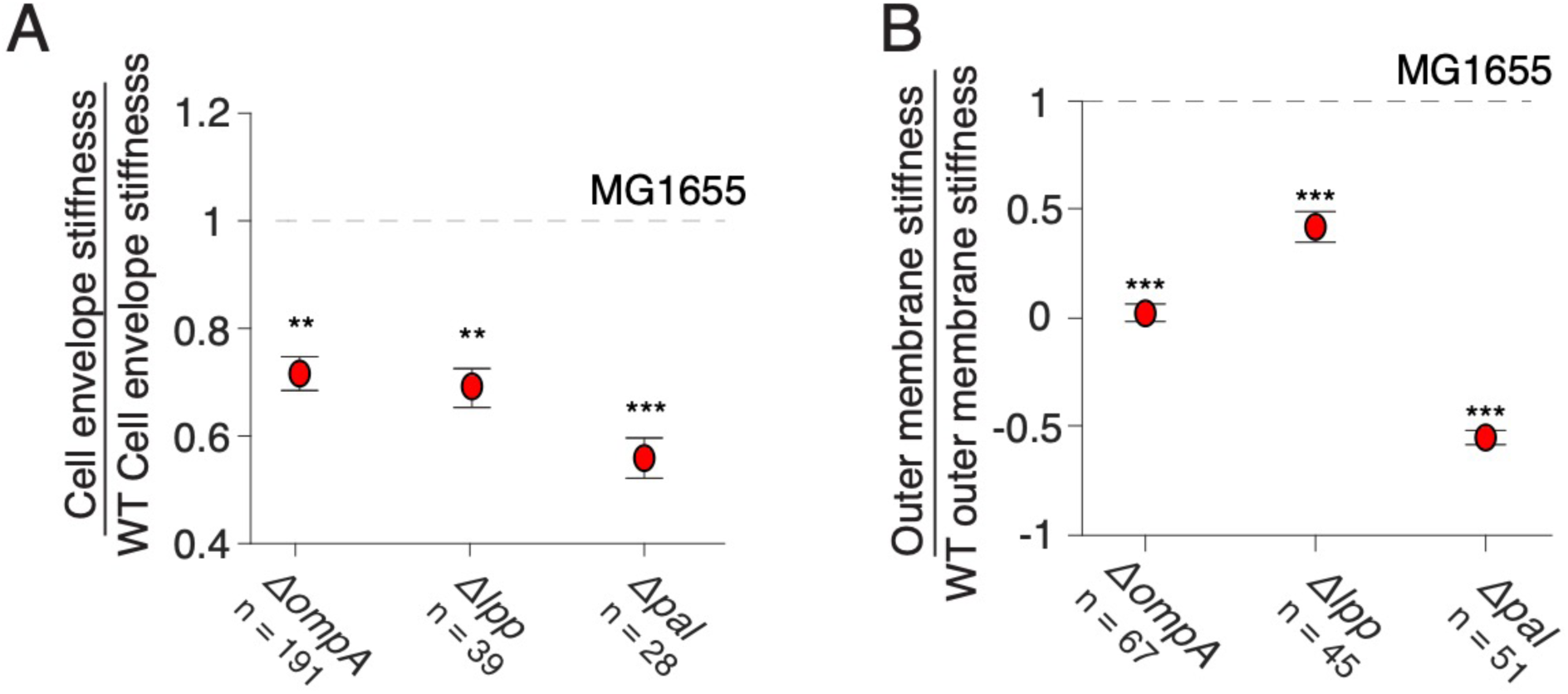
Deletion of Pal has a dramatic effect on cell envelope integrity. A) Cell envelope stiffness of mutants for outer membrane-cell wall linkers. B) Outer membrane stiffness of mutants for outer membrane-cell wall linkers.

**Figure 6.**
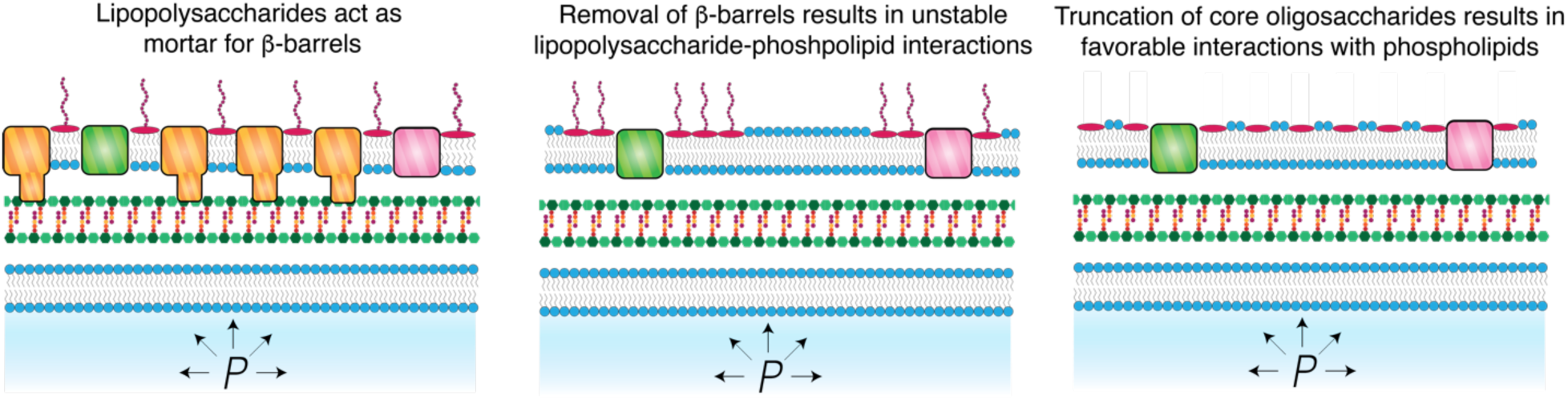
Model for the effect of the genetic interactions of *ompA* and *rfa* genes on cell envelope stiffness.

### Modifications to lipid A have weak effects on cell envelope stiffness

The above analysis demonstrates that hydrogen bonds between neutral sugar resides bear forces within the outer membrane. However, truncations to the core oligosaccharide require deletion of one of the *rfa* genes and are not viewed as adaptive except when cells are subjected to strong selective pressure such lytic bacteriophage predation^28^. Furthermore, most wild-type bacteria possess an O-antigen (which can also bear forces^1^), precluding phenotypic adaptation via modulation of core oligosaccharide length. On the other hand, it is well understood that bacteria use a suite of enzymes to adaptively modify lipid A in response to environmental cues^9^. By combinatorially expressing these enzymes this adaptive ability was previously exploited to synthetically engineer mutant *E. coli* strains that homogenously express precise variants of lipid A to investigate the dependence of the human immune response on lipid A chemistry^10^. For us, these mutants provided an opportunity to investigate the dependence of cell-envelope mechanics on lipid A chemistry and to explore whether, in principle, this chemistry could be used to mechanically adapt to their environment.

Our control strain (BN1) homogenously expressed hexaacylated, bis-phosphorylated lipid A (Fig. 1B), which is the most abundant species of lipid A in wild-type *E. coli*^10^. We hypothesized that reducing the negative charge of the head group would reduce outer membrane stiffness. Surprisingly, when we removed the 1-phosphate group, the stiffnesses of the total cell envelope was unaffected and the effective stiffness the outer membrane increased modestly (Fig. S2). Similarly, adding an acyl chain had little effect on cell envelope or outer membrane mechanical properties mechanics (Fig. S2).

## Discussion

Here, we developed a new quantitative assay to empirically calculate the stiffness of the bacterial cell envelope. In this assay, cells were subjected to a series of hyperosmotic shocks of increasing magnitude, and the contraction of cell envelope length was measured. A simple but important result from this experiment was that the degree of contraction upon each shock was linearly proportional to shock magnitude, which made it simple to unambiguously define envelope stiffness (Fig. 2D). This result is superficially at odds with previous atomic force microscopy-based measurements reporting non-linear mechanical properties of the cell wall^24^. However, AFM uses indentation to deform the cell envelope, causing stretching of the envelope rather than contraction. We anticipate that we would see similar non-linear strain-stiffening if we could controllably perform our assay using hypoosmotic shocks instead of hyperosmotic shocks. In fact, we previously noted that hypoosmotic shocks cause negligible swelling of the envelope of cells during steady-state growth. This could reflect extreme strain-stiffening, however it is difficult to control for the effect of stretch-activated ion channels^29^, which decrease pressure upon hypoosmotic shocks and would therefore reduce cell envelope swelling. Furthermore, deletion of channels causes cell lysis upon hypoosmotic shock, making the control experiment impossible. We simply conclude that the cell envelope is linearly elastic for pressures up to the steady-state pressure at which cells grow. Interestingly, our initial applications of the osmotic force extension assay to Gram-positive bacteria reveal that it is linear elastic over a wide range of positive and negative pressure variation.

A second central finding of our study is that the core oligosaccharide moieties of lipopolysaccharides only contribute to cell envelope stiffness if the outer membrane possesses its full complement of β-barrel proteins. These proteins completely pack the outer membrane and in this light, lipopolysaccharides function as “mortar” that fills the space between β-barrels (Fig. 5A). The deletion of OmpA, one of the most abundant β-barrel proteins, leaves a void in the outer membrane filled by phospholipids in both the inner and outer leaflets of the outer membrane. Furthermore, in this mutant, lipids and proteins phase separate from the β-barrels proteins in the outer membrane ^13^. The specific spatial pattern of phospholipids and lipopolysaccharides in the lipid phase is unknown, but based on our results we hypothesize that lipopolysaccharides and phospholipids further phase separate due to self-affinity, for example, between core oligosaccharides (Fig. 5B). This would be expected to lead to a fragile outer membrane due to line tension at the boundary of phospholipid-lipopolysaccharide domains. Therefore, truncation of core oligosaccharides in the *ΔompA* background reduces self-affinity of lipopolysaccharides, leading to mixing that increases the stiffness of the outer membrane (Fig. 5C). It is not possible to directly image lipopolysaccharides, but this hypothesis will be interesting to test in future studies.

Our model is consistent with a model that was recently proposed to explain the mechanical phenotype of the *ΔbamD* mutant^30^. BamD is a regulatory lipoprotein that activates the outer membrane Bam complex^31^, which folds β-barrel proteins into the outer membrane. Deletion of *bamD* globally reduces β-barrel content in the outer membrane and, like the deletion of *ompA*, results in outer leaflet phospholipids. It was proposed that this leads to tension in the outer membrane, which renders the cell fragile to osmotic fluctuations. This was supported by the finding that inhibiting constitutive removal of phospholipids from the outer leaflet by the Mla and PldA systems increases cell viability during fluctuations. Based on our data, we hypothesize that truncating the core oligosaccharide in the *ΔbamD* would also have a protective effect.

An interesting observation from the sum of our measurements is that full truncation of core oligosaccharides, deletion of OmpA, and deletion of other outer membrane-cell wall linkers all caused the same quantitative reduction in total cell envelope stiffness (≈20-25%; Fig. 3A,4A,5A). We propose that this convergent phenotype points to a common structural cause for envelope weakening: minor delamination (but not complete detachment) of the outer membrane from the cell wall. In this model stiffness of the outer membrane plays two related roles: i) it bears in-plane compression and ii) it prevents out-of-plane buckling, which limits its ability to bear in-plane compression geometrically. This is consistent with our observation that expression of the OmpA β-barrel alone, without the periplasmic linker domain, is sufficient to prevent bulging of the outer membrane upon hyperosmotic shock (Fig. 4C,D). It will likely be possible to test this model explicitly by combining the osmotic force extension assay with super-resolution measurements of outer membrane geometry.

Contrary to total envelope stiffness, various perturbations to outer membrane composition had a wide range of effects on outer membrane stiffness (Fig. 3B, 4B, 5B), when this quantity was decoupled from total envelope stiffness using the plasmolysis-lysis assay. This likely means that these mutations differentially affect the outer membrane’s ability to stay mechanically engaged to the cell wall for large hyperosmotic shocks.

Our most surprising result was that modification to lipid A – including those to the head group and the acyl chains - had no effect on outer membrane mechanics. Interestingly, this means that divalent cation-mediated bridging of adjacent lipopolysaccharides molecules has a much greater effect on outer membrane permeability than mechanics.

Collectively, our analysis suggests that the global mechanical properties of the cell envelope arise from complex interactions between the various components of the envelope, rather than additive contributions from each component.

## Author contributions

DF conceptualized the study, acquired funding, performed microfluidic assays and MD simulations, analyzed data, generated bacterial strains, and wrote the manuscript. AA and TS performed microfluidic assays, analysis, and bacterial strain generation. GH conceptualized the study and acquired funding. ERR conceptualized the study, acquired funding, and wrote the manuscript. All authors contributed to discussing the data. DF, GH and ERR reviewed and edited the manuscript.

## Acknowledgement

We thank Steven Trent and Carmen Herrera for providing us with bacterial strains. We thank Georgina Benn for helpful discussion. DF was funded by an NSF Graduate Research Fellowship. ERR was supported by NIH Grant R35GM143057. GMH was supported by NIH Grant R35GM138312.

## Methods

### Bacterial strains and culture conditions

Bacterial strains and plasmids used in this study are listed in Table S1. Bacteria were grown in lysogeny broth (LB), Lennox formulation (5 g l^-1^ NaCl) overnight in a rotary shaker at 37°C. For selection, 50 µg/ml kanamycin or 100 µg/ml ampicillin were used. The osmolarity of the growth medium was modulated with sorbitol (Sigma).

### Construction of chromosomal gene deletion mutants

P1 vir phage transduction was used to move selectable deleted genes from the donor BW25113 strain to the recipient MG1655 strain^32^. Mutations were confirmed by PCR using primers that anneal outside of flanking regions of deleted gene (Table S2). When necessary, excision of the resistance gene was carried out using the helper plasmid pCP20^33^.

### Lambda phage recombineering

To generate the mutant allele of *ompA* lacking the periplasmic domain (*11ompA^PD^*) mutant we used lambda red recombineering. Cells carrying the red recombinase expression plasmid, pKD46, were grown in 30mL LB with ampicillin at 30°C to an OD of 0.4. The culture was then inoculated with L-arabinose to final concentration of 10% and incubated at 30°C for an additional 15 min. To make electrocompetent cells, the culture was initially chilled on ice before undergoing two washes with cold ultrapure deionized water, and a final wash with ice-cold 10% glycerol in water.

Electrocompetent cells were aliquoted into 30μL suspensions and stored at −80°C. Electroporation was conducted by using a Gene Pulser Xcell Electroporator with 0.2cm electrode gap cuvettes. 30 μL of competent cells were inoculated with at least 200ng of DNA, and shocked with a 2.5kV voltage. Shocked cells were immediately recovered with 1mL SOC media and incubated with shaking at 30°C for two hours. Next, 500μL of transformant culture was spun down, resuspended in 100μL LB, and spread onto agar plates to select for kanamycin-resistant transformants. Plates were left growing overnight at 37°C.

The kanamycin cassette, which included FRT sites, was first amplified from pKD13 using Primer 1 and Primer 2 (Table S1). Primer 1 contains a 50bp homologous region to the 3’ end of the *ompA β*-barrel domain, in addition to the first 20bp of the 5’ region of the resistance marker. Primer 2 contains a 50bp homologous region downstream of the stop codon of *ompA*, and the last 20bp of the 3’ end of the resistance marker. After confirmation of a 1.4kp amplicon, the remaining PCR product was treated with DpnI for two hours at 37°C, followed by column purification. The purified, linear DNA was used for electroporation of BW25113 cells carrying the red recombinase expression plasmid, pKD46, following the protocol described above. After primary selection, P1 vir phage transduction was used to move the truncated *ompA* gene with kanamycin resistance into the recipient MG1655 background. Chromosomal integration of *△ompA^PD^::kan* was verified through colony PCR using primer pairs TS023/TS024 and AA001/AA002.

### Imaging in microfluidic devices

Cells were imaged on a Nikon Eclipse Ti-E inverted fluorescence microscope with a 100X (NA 1.45) oil-immersion objective. For all experiments we used CellASIC B04A microfluidic perfusion plates and medium was exchanged using the CellASIC ONIX microfluidic platform. Images were collected on a sCMOS camera (Prime BSI). Experiments were performed at 37°C in a controlled environmental chamber (HaisonTech).

### Combinatorial color-coding

To accelerate our screen for the effect of genetic perturbations on cell envelope stiffness, we typically measured three strains at a time by color-coding them with non-toxic dyes, pooled the three strains, performed experiments on the pool, and then decoded our color-code using custom computational image analysis.

For color-coding, bacterial plasma membranes and/or cell walls were stained either individually or in two-color combination. Controls were performed to ensure measurements were not affected by dyes, and unless otherwise stated, each experiment was repeated in triplicate whereby in each experiment the color-code was permuted across the three strains. We included a wild-type control in each set of three mutants. To color code, the cell envelope was stained with wheat germ agglutinin–AlexaFluor488 (WGA-AF488, Life Technologies), the fluorescent D-amino acid HADA (Tocris Bioscience), MitoTracker Orange CM-H_2_TMRos (Invitrogen), MitoView Green (Biotium), MitoView 650 (Biotium), MitoView 720 (Biotium). MitoTracker Orange CM-H_2_TMRos (250 nM), MitoView Green (200 nM), or MitoView 720 (100 nM) were added to diluted cultures 30 minutes before pooling strains. HADA (250µM) was added to diluted cultures 1 hour before pooling.

### Osmotic force extension assay

Overnight cultures were diluted 100-fold into 1 ml of fresh LB and incubated for 2 h with shaking at 37 °C. Plates were loaded with medium pre-warmed to 37 °C. 5μg /mL Alexa Fluor 647 NHS Ester dye was added to specific media as a tracer dye for medium switching.

Cells were grown for 5 min in LB in the imaging chamber before being subjected to a series of hyperosmotic shocks using LB with 50mM, 100 mM, 200 mM, and 400 mM sorbitol for 1 minute each. Between sorbitol shocks the media was switched back to LB for 1 minute.

To calculate the amplitude of length oscillations during osmotic shocks, cells were tracked using custom MATLAB algorithms. First, cell-envelope lengths (*l*) were automatically detected and the elongation rate 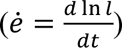 was calculated for each cell. The effective population-averaged length was calculated by integrating the population-averaged elongation rate over time^34^. The mechanical strain in cell envelope length caused by each hyperosmotic shock 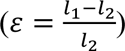 was then calculated. Linear regression of mechanical strain as a function of shock magnitude was calculated where cell-envelope stiffness was defined as the inverse of the slope of the regression. Uncertainty was estimated using the standard error of the linear regression.

To control for experiment-to-experiments variability due to heterogeneity in microfluidic chips, e normalized cell-envelope stiffness to the internal wild-type control in each experiment before averaging across experiments.

### Plasmolysis-lysis experiments

Plasmolysis-lysis experiments were performed as described previously^1^, with minor modifications. Briefly, overnight cultures were diluted 100-fold into 1mL of fresh LB media and incubated with shaking at 37°C for one hour. 250 µM of HADA was added to the culture and cells were incubated for an additional hour. Cultures were then back diluted 100-fold into 1mL of pre-warmed LB with 250 µM of HADA, which we added directly to the loading well of the microfluidic chip. After loading cells into the imaging chamber they were perfused with LB for 5 min, followed by LB + 3M sorbitol for 5 min, then with LB + 3M sorbitol + 20% N-lauroylsarcosine sodium salt (Sigma) for 30 min, and finally with LB for 20 min. We measured the cell wall length upon lysis after this last step to control for the possibility that the detergent had direct effects on cell wall rest length. 1uL of Alexa Fluor 647 NHS Ester dye (1mg/mL) was added to every other perfusion well as a tracer dye to track media switching. A time-lapse image with a 10 s frame rate was taken during the initial 5 min period when the cells were perfused with LB. To avoid photobleaching of HADA and phototoxicity, a single image was taken during each of the next two perfusion periods when the cells were plasmolyzed (LB+ 3M sorbitol) and detergent-lysed (LB+ 3M sorbitol+ N-lauroylsarcosine sodium salt), respectively.

As before, the outer membrane and the cell wall were treated as parallel linear springs and the relative stiffnesses were calculated as:

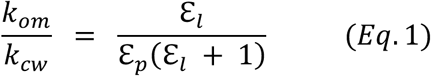

where ℇ_p_ is the strain induced in the cell wall upon plasmolysis with 3 M sorbitol and ℇ_1_ is the additional strain induced by the detergent lysis of the cell (Fig. S1). By further substituting the total envelope stiffness (*k*_env_=*k*_cw_+*k*_om_) into **Eq. 1** the stiffnesses of the cell wall and outer membrane were explicitly solved for in terms of experimentally measurable quantities:

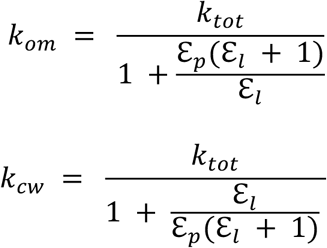

### Outer membrane bulging experiments

For cell bulging experiments, the outer membrane was labeled with WGA-AF488, which was added to the loading well to a final concentration of 10 ng ml−1.

### Molecular Dynamic Simulations

*E. coli* (K12) outer membrane models were built with five distinct lipopolysaccharide cores corresponding to the forms produced by *△rfaC*, *△rfaF*, *△rfaG*, *△rfaJ*, and WT (Fig. 1B), using the CHARMM-GUI online server^35^ with CHARMM36 force field parameters^36, 37^ and TIP3P water. Lipopolysaccharide bilayers were generated to probe only the contribution of this molecule to the outer membrane. Simulated bilayers contained 53 LPS molecules on both the outer and inner leaflets. The minimum water height on the top and bottom of the system was set to 40 Å. Systems were minimized and equilibrated using the CHARMM-GUI lipids protocol.^35^ Production simulations were performed at 310.15K in NPT.

Production data were collected using GROMACS 2020.4 molecular dynamics (MD) engine^38^ patched with PLUMED version 2.7.0^39^. Lipopolysaccharide bilayers were run for 300 ns using a 2 fs timestep at lateral pressure *P* = 0 for an initial equilibration after which the pressure was changed to either 10, 25, 50, 100 bar, and run for an additional 300 ns.

---

**Figure S1.**
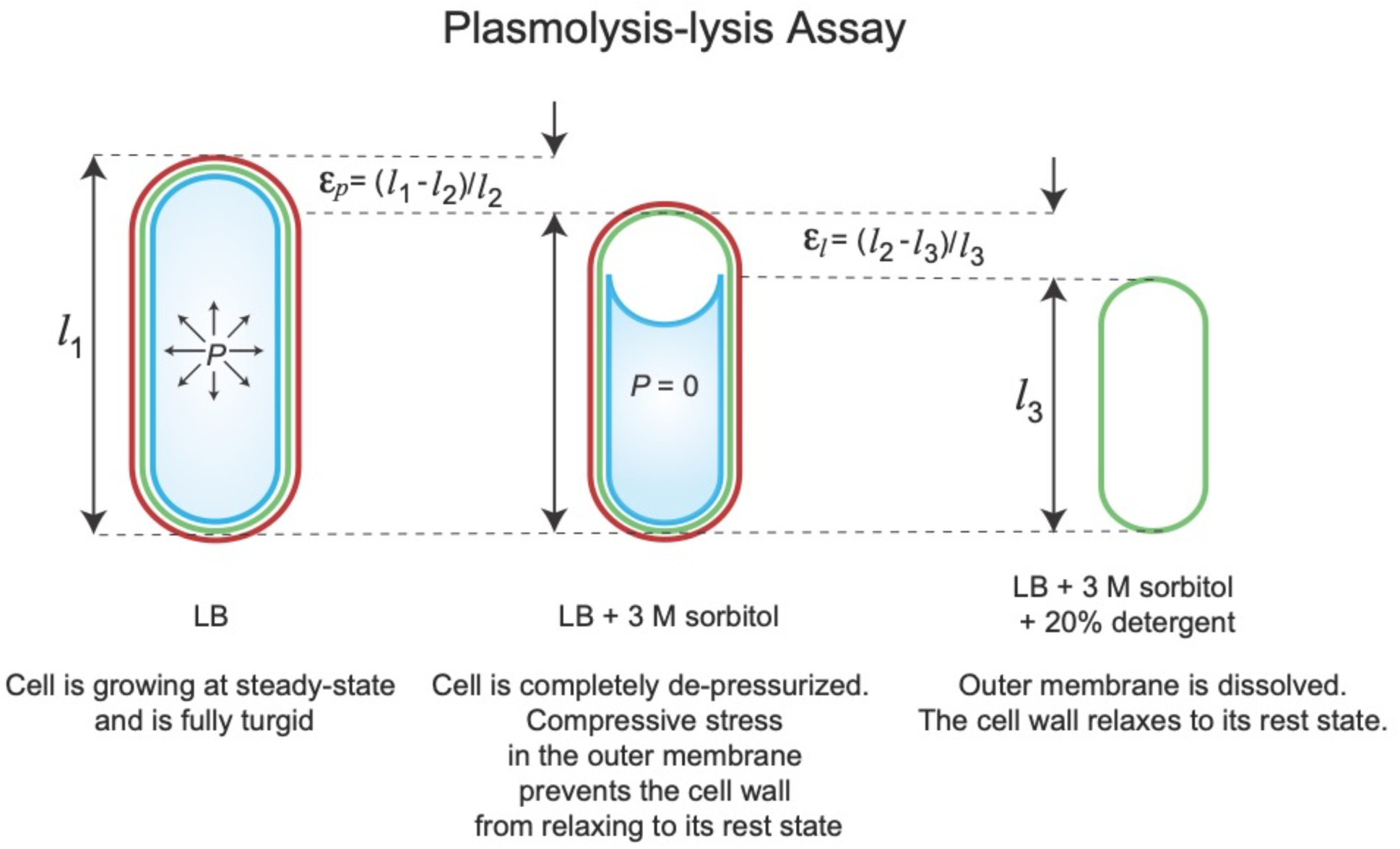
Plasmolysis-lysis assay used to measure the ratio between the stiffness of the cell wall and the outer membrane. Model of a fully turgid cell at a steady-state length (*l_1_*). The cell is de-pressurized by a large 3M hypo-osmotic shock resulting in a plasmolysed cell whose length contracts (*l_2_*). The strain resulting from this shock is calculated by: 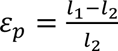. The cell is then treated with 20% detergent which dissolves the outer membrane allowing the cell wall to relax to its rest state (*l_3_*). The strain resulting from this shock is calculated by: 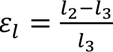. By treating the outer membrane and cell wall as parallel linear springs, relative stiffness is calculated by: 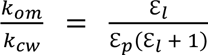.

**Figure S2.**
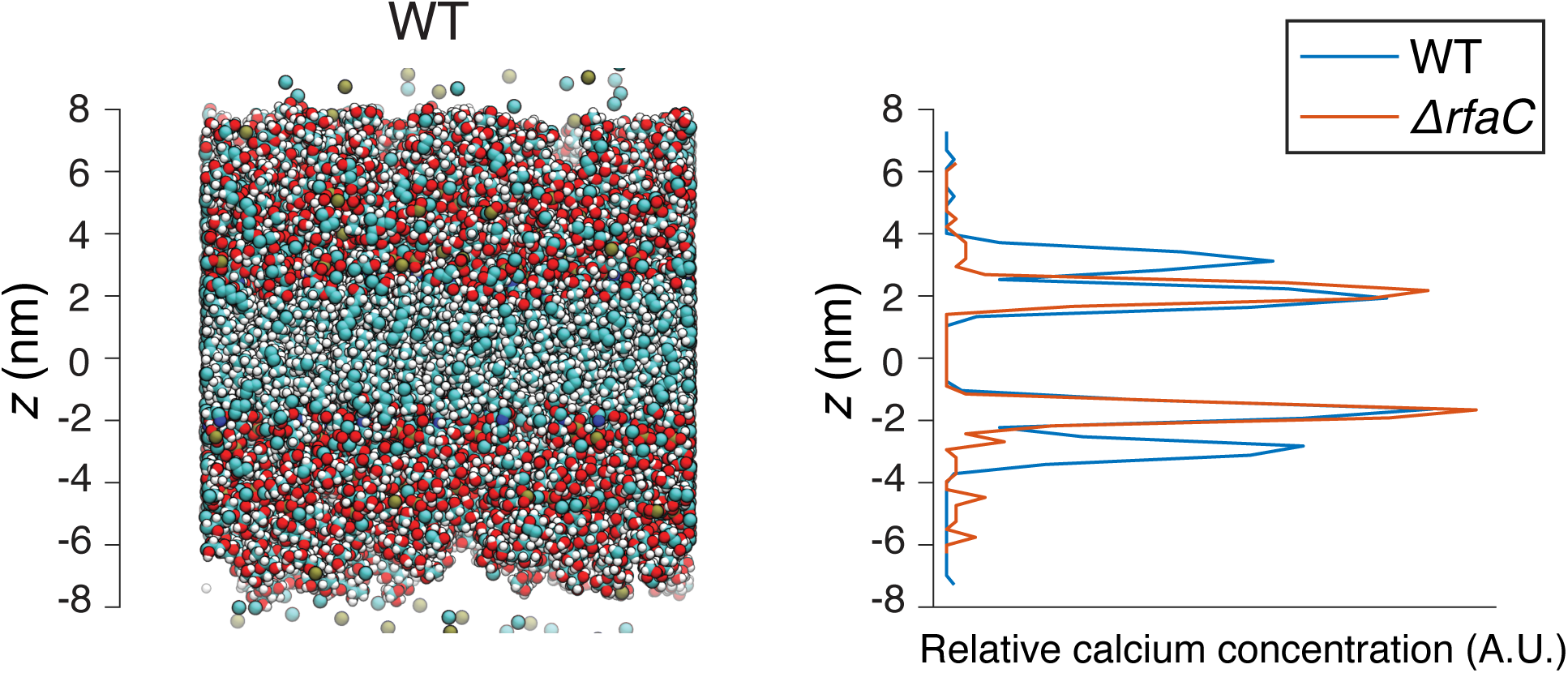
Calcium distribution reflects the phosphate distribution in simulated lipopolysaccharide bilayers. A) Simulated wild-type lipopolysaccharide bilayer. B) Calcium distribution across the thickness of the bilayer.

**Figure S3.**
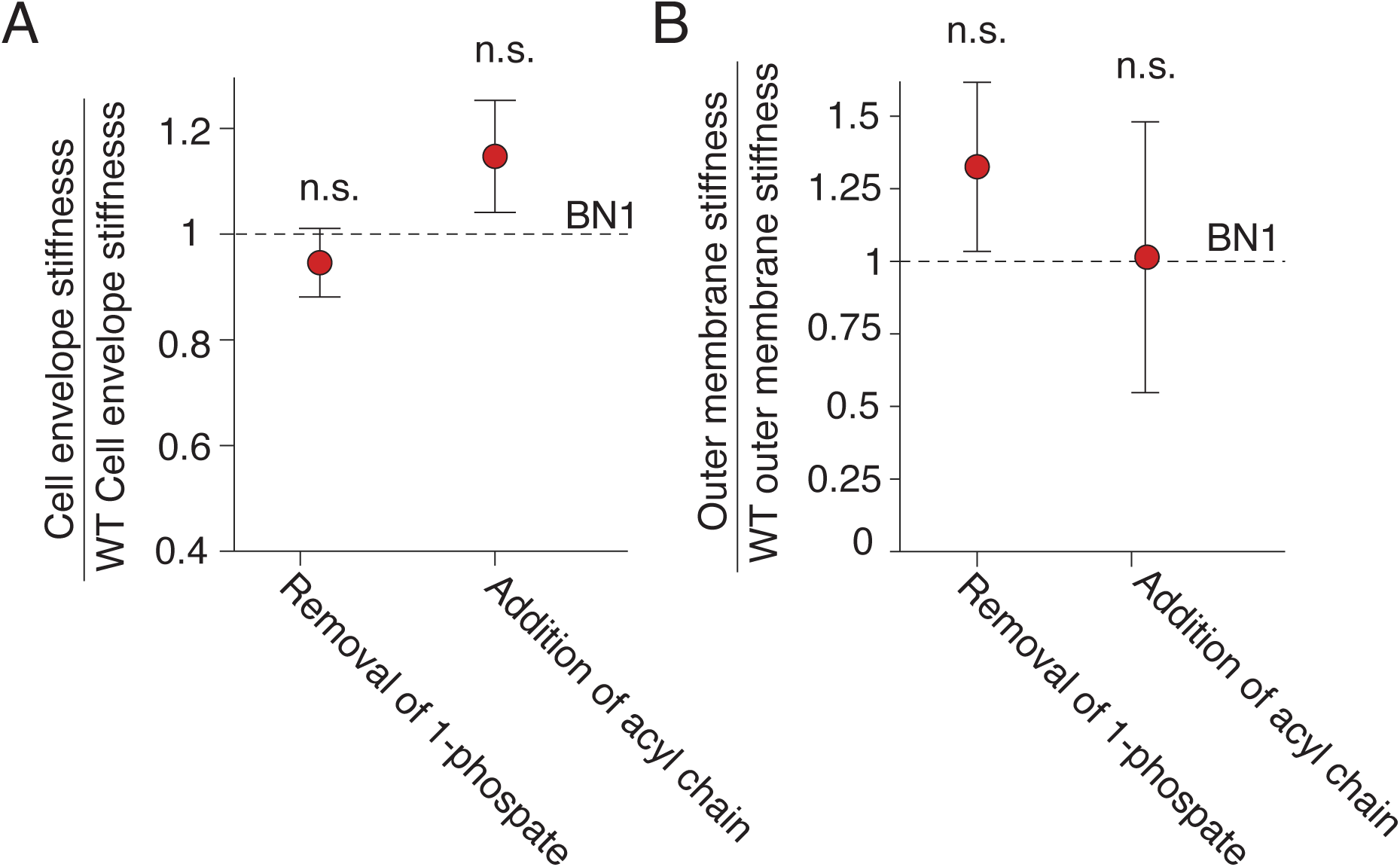
Modifications to lipid A have weak effects on cell envelope stiffness. A) Cell envelope stiffness of modified lipid A strains, normalized by wild-type (BN1) cell-envelope stiffness; n = 48, 54, 64, for BN1pE (removal of 1-phosphate), BN1pP (addition of acyl chain), and BN1 wild-type cells. B). Outer membrane stiffness of normalized by wild-type (BN1) cell-envelope stiffness; n = 45, 51, 87, for BN1pE, BN1pP, and BN1 wild-type cells.

**Table S1.**
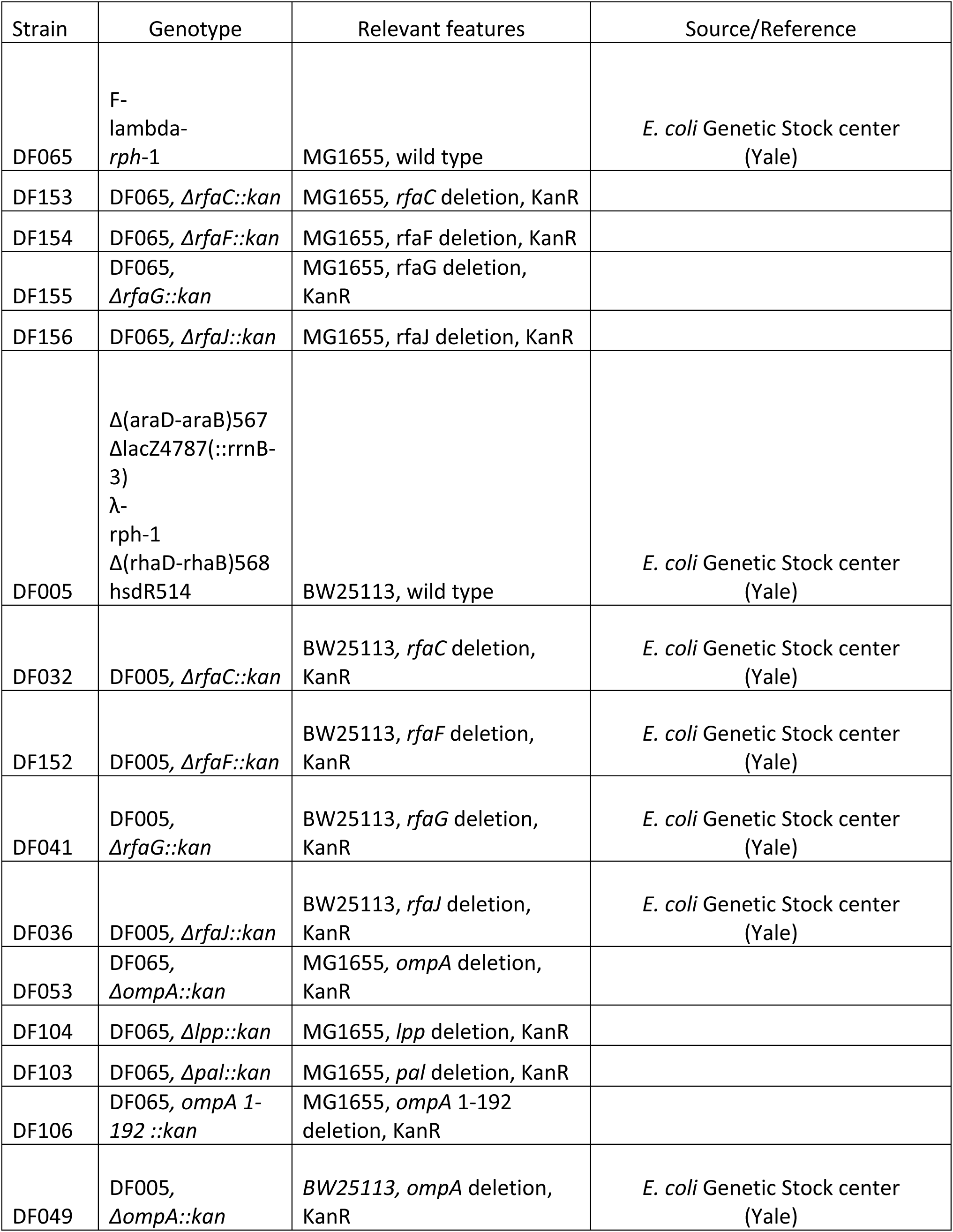
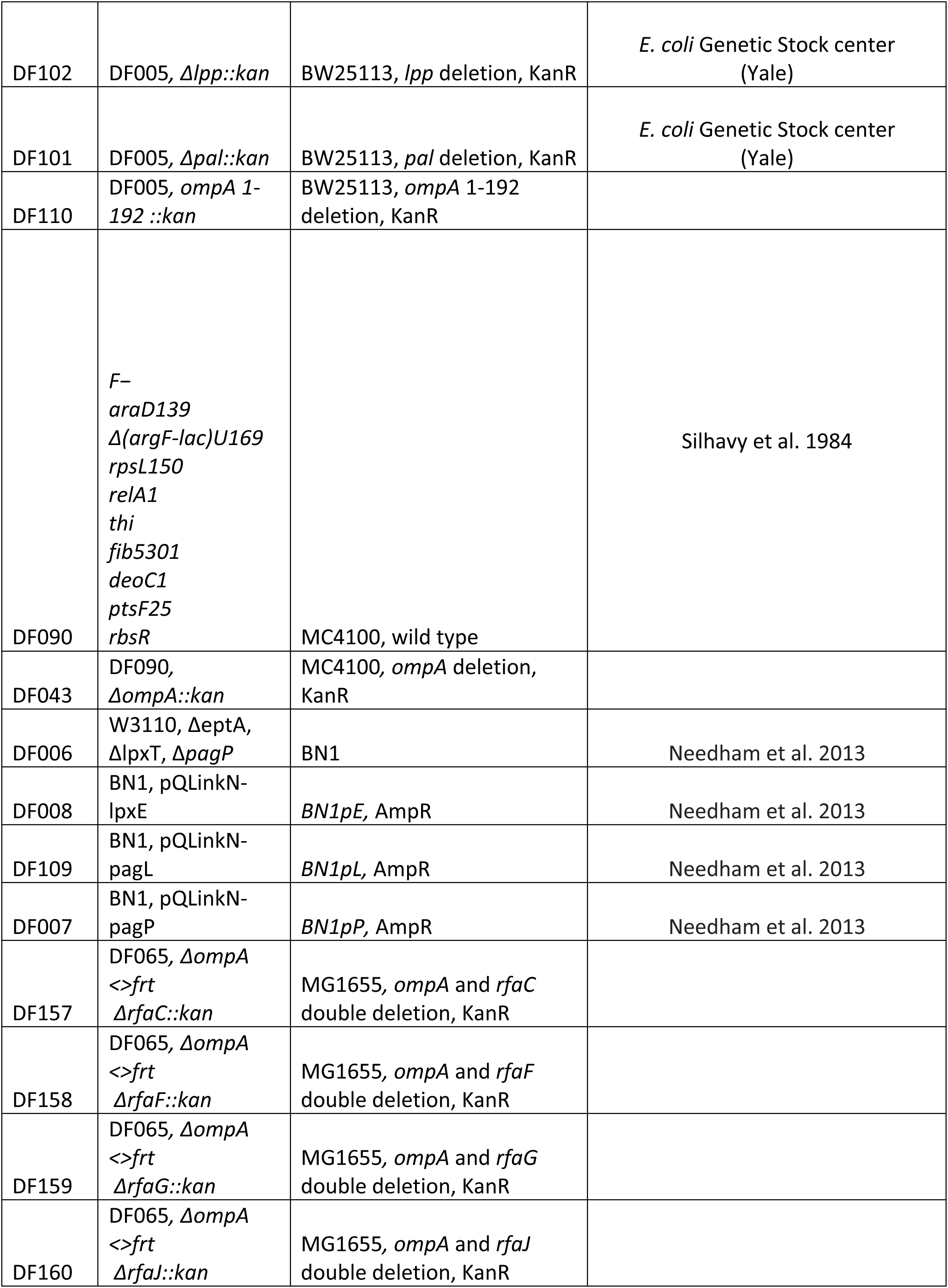
Strains used in this study.

**Table S2.**
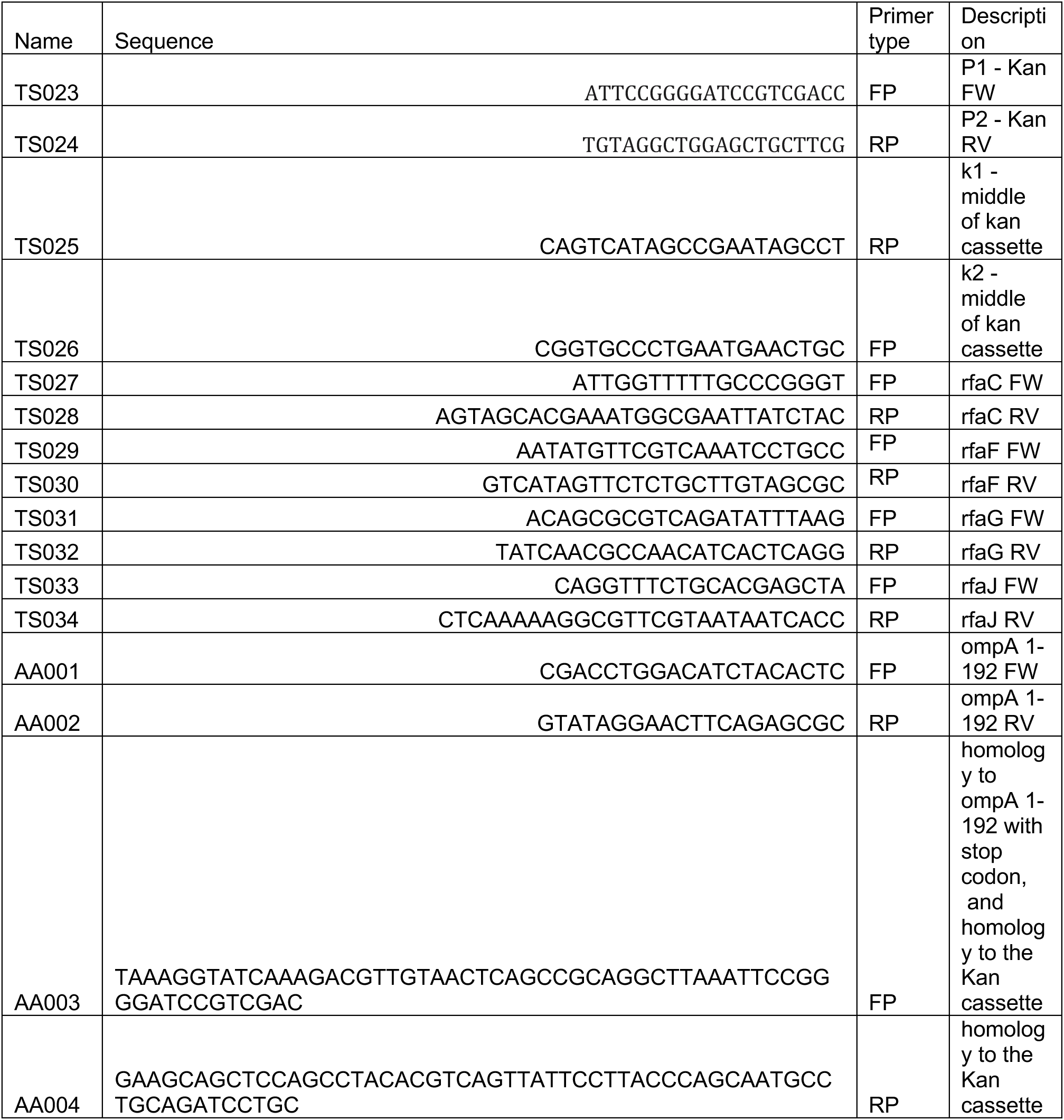
Primers used in this study.

